# Ectopic insert-dependent neuronal expression of GFAP promoter-driven AAV constructs in adult mouse retina

**DOI:** 10.1101/2022.04.06.487191

**Authors:** Nguyet Le, Haley Appel, Nicole Pannullo, Thanh Hoang, Seth Blackshaw

## Abstract

Direct reprogramming of retinal Müller glia is a promising avenue for replacing photoreceptors and retinal ganglion cells lost to retinal dystrophies. However, questions have recently been raised about the accuracy of studies claiming efficient glia-to-neuron reprogramming in retina that were conducted using GFAP mini promoter-driven adeno-associated virus (AAV) vectors. In this study, we have addressed these questions using GFAP mini promoter-driven AAV constructs to simultaneously overexpress the mCherry reporter and candidate transcription factors predicted to induce glia-to-neuron conversion, in combination with prospective genetic labeling of retinal Müller glia using inducible Cre-dependent GFP reporters. We find that, while control GFAP-mCherry constructs express faithfully in Müller glia, 5 out of 7 transcription factor overexpression constructs tested are predominantly expressed in amacrine and retinal ganglion cells. However, genetic cell lineage analysis shows no evidence for glia-to-neuron conversion. These findings demonstrate strong insert-dependent effects on AAV-based GFAP mini promoter specificity that preclude its use in inferring cell lineage relationships when studying glia-to-neuron conversion in retina.

## Introduction

Retinal degeneration is characterized by irreversible loss of retinal neurons and is a major cause of blindness. Macular degeneration and retinitis pigmentosa induce rod and cone photoreceptor loss, while glaucoma leads to retinal ganglion cell degeneration. Two main classes of cell-based therapies are being actively pursued to replace these lost neurons. The first of these involves direct differentiation and transplantation of ES/iPS-derived cells, for which large numbers of the desired cell type can be produced on demand (Osakada et al., 2008; Lamba et al., 2010; Tucker et al., 2013). However, this approach has thus far had limited success, largely because of the limited efficiency of integration of transplanted cells into host circuitry, particularly in diseased retina (Gasparini et al., 2019; Miltner and La Torre, 2019; Singh et al., 2020; Zhang and Wang, 2022). The second approach takes inspiration from injury-induced regeneration that occurs in many cold-blooded vertebrates, and involves directed reprogramming of retinal Müller cells into neurogenic progenitors and/or retinal neurons (Goldman, 2014; Lahne et al., 2020; Salman et al., 2021). Glial reprogramming has a number of potential therapeutic advantages relative to exogenous cell transplantation, including the reduced likelihood of immune rejection and the fact that glial-derived neurons are already present in the retina itself.

Studies using transgenic mice for cell lineage tracing have shown that either overexpression of the neurogenic bHLH factor *Ascl1* (Jorstad et al., 2017; Todd et al., 2020, 2021), or genetic disruption of NFI family transcription factors (Hoang et al., 2020) induces Müller glia to generate bipolar and amacrine interneurons, but neither photoreceptors nor mature retinal ganglion cells. In contrast, several studies using adeno-associated virus (AAV)-mediated transcription factor overexpression (Yao et al., 2018; Xiao et al., 2021) or CasRx-mediated knockdown of the splicing regulator *Ptbp1* (Zhou et al., 2020) have reported highly efficient generation of photoreceptor and retinal ganglion cells from Müller glia.

Particularly in the light of similar studies reporting efficient astrocyte-to-neuron reprogramming in brain (Brulet et al., 2017; Qian et al., 2020; Zhou et al., 2020; Xiang et al., 2021), AAV-based glia-to-neuron conversion has been hailed as a broadly promising approach for treatment of many classes of neurodegenerative disorders (Janowska et al., 2019; Götz and Bocchi, 2021; Wang et al., 2021b). However, several important technical concerns have arisen about many of these studies. First, while studies reporting Müller glia-to-neuron conversion using transgenic mice have used well-characterized inducible transgenic Cre lines to infer cell lineage relationships (Jorstad et al., 2017; Hoang et al., 2020), AAV-based studies have used GFAP mini promoter-driven Cre constructs for this purpose. Second, the GFAP minipromoter, which is used to drive glial-specific expression of constructs used for reprogramming, can show leaky neuronal expression in brain (Taschenberger et al., 2017; Griffin et al., 2019), most notably when overexpressing *Neurod1* or disrupting *Ptbp1* expression in brain astrocytes (Chen et al., 2021; Wang et al., 2021a), both of which have been claimed to induce astrocyte-to-neuron conversion (Puls et al., 2020; Qian et al., 2020; Zhou et al., 2020; Xiang et al., 2021).This raises the question of whether reports of AAV-based Müller glia-to-neuron conversion actually represent cases of ectopic expression of GFAP reporter constructs in endogenous neurons (Blackshaw and Sanes, 2021; Qian et al., 2021).

To directly address this question, we used tamoxifen treatment to selectively label Müller glia in adult mouse retina using the well-characterized *GlastCreERT2;Sun1-GFP* transgenic line (de Melo et al., 2012; Hoang et al., 2020). We then performed intravitreal injection in these retinas with GFAP promoter-driven 7m8 AAV2 constructs expressing either mCherry alone or mCherry in combination with a range of different transcription factors predicted to potentially induce glia-to-neuron conversion. We found that Sun1-GFP expression was restricted to Müller glia, but that a number of constructs overexpressing transcription factors showed leaky mCherry expression in multiple inner retinal neuronal subtypes, including retinal ganglion cells. This demonstrates widespread insert-dependent induction of leaky neuronal expression of GFAP minipromoter-driven constructs and underlines the importance of using well-characterized genetic cell lineage analysis when studying glia-to-neuron conversion.

## Results

We sought to measure Müller glia-to-neuron conversion following overexpressing of candidate transcription factors using GFAP minipromoter-based 7m8 AAV2 vectors in the adult mouse retina (Aartsen et al., 2010; Hickey et al., 2017; Yao et al., 2018; Zhou et al., 2020). Constructs tested included a GFAP-mCherry control and a range of different transcription factors that have been previously reported to reprogram Müller glia or were candidates for doing so based on their role in regulating neurogenesis and/or retinal neuron specification (Fig. 1A) (Mears et al., 2001; Sharma et al., 2019; Puls et al., 2020; Lyu et al., 2021; Todd et al., 2021; Xiao et al., 2021). To genetically label Müller glia prior to AAV infection, we conducted 3 consecutive intraperitoneal injections of 4-hydroxytamoxifen (4-OHT) from P21-P23 in *GlastCreER;Sun1-GFP* mice (Fig. 1A). At 2 months of age, we then performed intravitreal AAV injection, and analyzed retinas at 8 and 21 days later (Fig. 1A). The *GlastCreER;Sun1-GFP* mice show selective and robust perinuclear GFP labeling which colocalizes with selective Müller glial marker Sox9 (Fig. 1B).

**Figure 1:**
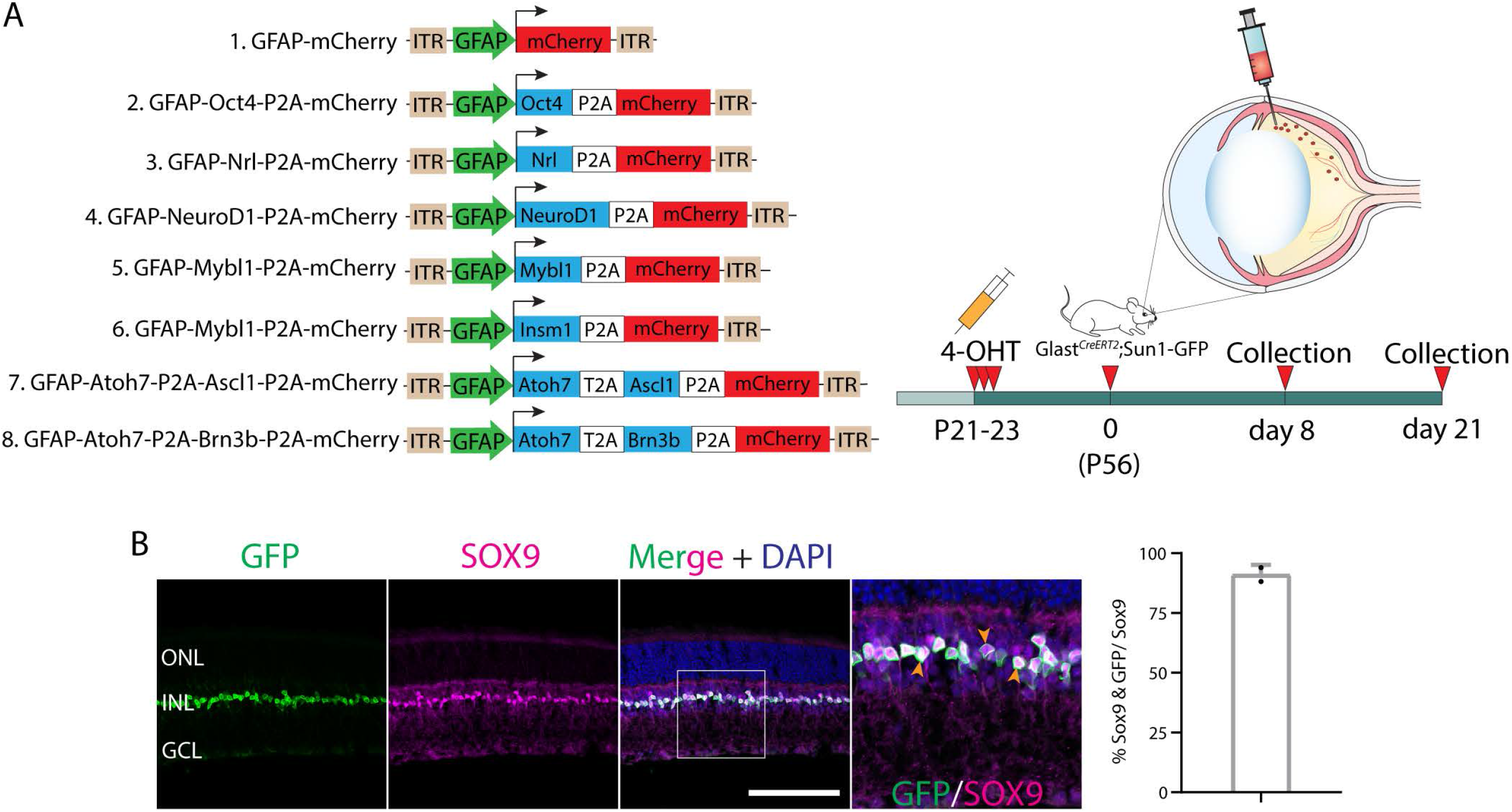
(A) Schematic of the GFAP AAV constructs and the workflow: three weeks old *GLASTCreERT2;Sun1-GFP* mice were induced with 3 daily doses of 4-OHT delivered intraperitoneally, intravitreally injected with GFAP AAV constructs at 2 months of age, and collected 8 days and 21 days following AAV infection for analysis. **(B)** Representative immunostaining for GFP and Sox9 expression in retinas following GFAP AAV infection. Quantification of the number GFP+ cells co-labeled with Sox9+ cells showed robust and selective expression in Sun1-GFP and Sox9-positive Müller glia. Yellow arrowheads indicate GFP+/Sox9+ cells. ONL, outer nuclear layer; INL, inner nuclear layer; GCL, ganglion cell layer. Scale bar = 100μm.

At both 8 and 21 days after AAV injection, we saw considerable differences in the pattern of cell type-specific expression of mCherry among the different constructs tested. GFAP-mCherry control, GFAP-Oct4-mCherry, and GFAP-Nrl-mCherry all showed almost complete colocalization of mCherry expression with Müller glia-specific Sun1-GFP (Fig. 2A-C, 3A-C, 4A-C). In these constructs, little to no mCherry expression overlapped with the amacrine cell-specific marker Tfap2a (Fig. 2), the ganglion cell-specific marker Rbpms (Fig. 3), or the bipolar/photoreceptor cell-specific marker Otx2 (Fig. 4) (Lyu et al., 2021). All other constructs tested – including GFAP-Neurod1-mCherry, GFAP-Mybl1-mCherry, GFAP-Insm1-mCherry, GFAP-Atoh7-Ascl1-mCherry, and GFAP-Atoh7-Brn3b-mCherry – showed little mCherry expression in Sun1-GFP-positive Müller glia (Fig. 2D-I, 3D-I, 4D-I). Instead, robust mCherry expression was detected in the inner retinal neurons, specifically retinal amacrine cells (Fig. 2J) and ganglion cells (Fig. 3J), although no mCherry expression was seen in Otx2-positive retinal bipolar cells for any of the constructs tested (Fig. 4). This pattern of retinal neuronal expression was also evident at 8 days following infection for GFAP-Insm1-mCherry, GFAP-Atoh7-Ascl1-mCherry, and GFAP-Atoh7-Brn3b-mCherry (Fig. S1-S3).

**Figure 2:**
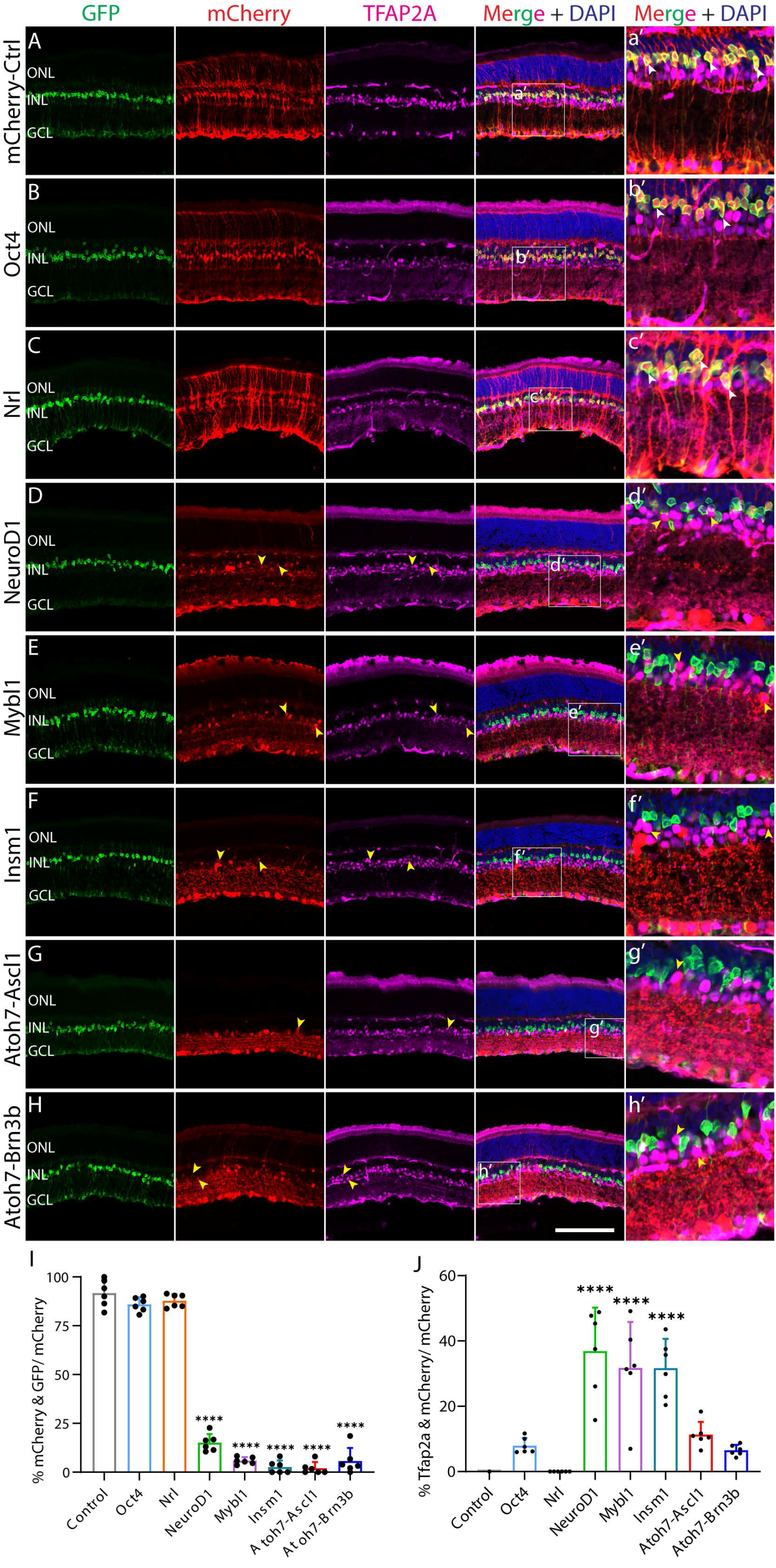
GFAP promoter constructs show insert-dependent lack of Müller glia-specific expression and ectopic mCherry expression in amacrine cells in adult mouse retina. **(A-H)** Representative immunostaining for GFP, mCherry and Tfap2a expression in the retinas collected 21 days post GFAP AAV infection. **(a’-h’)** Outlined region shown at higher magnification. GFAP-mCherry **(A, a’)**, GFAP-Oct4-mCherry **(B, b’)** and GFAP-Nrl-mCherry **(C, c’)** showed almost complete colocalization of construct-derived mCherry and Müller glia-specific Sun1-GFP. White arrowheads indicate co-labeled mCherry+/GFP+ cells. GFAP-NeuroD1-mCherry **(D, c’)**, GFAP-Mybl1-mCherry **(E, e’)** and GFAP-Insm1-mCherry **(F, f’)** GFAP-Atoh7-Ascl1-mCherry **(G, g’)**, and GFAP-Atoh7-Brn3b-mCherry **(H, h’)** showed low mCherry/GFP colocalization. Robust mCherry expression in amacrine cells is observed in these GFAP AAV constructs **(D, d’-F, f’)**. Yellow arrowheads indicate Tfap2a+/mCherry+ double positive cells. Quantification of mean percentage ± SD of mCherry+/GFP+ cells **(I)** and Tfap2a+/mCherry+ cells **(J)**. Significance was determined via one-way ANOVA with Dunnett’s test: ***p < 0.001, ****p < 0.0001. Each data point in the bar graphs was calculated from an individual retina (*n* = 6). ONL, outer nuclear layer; INL, inner nuclear layer; GCL, ganglion cell layer. Scale bar = 100μm

**Figure 3:**
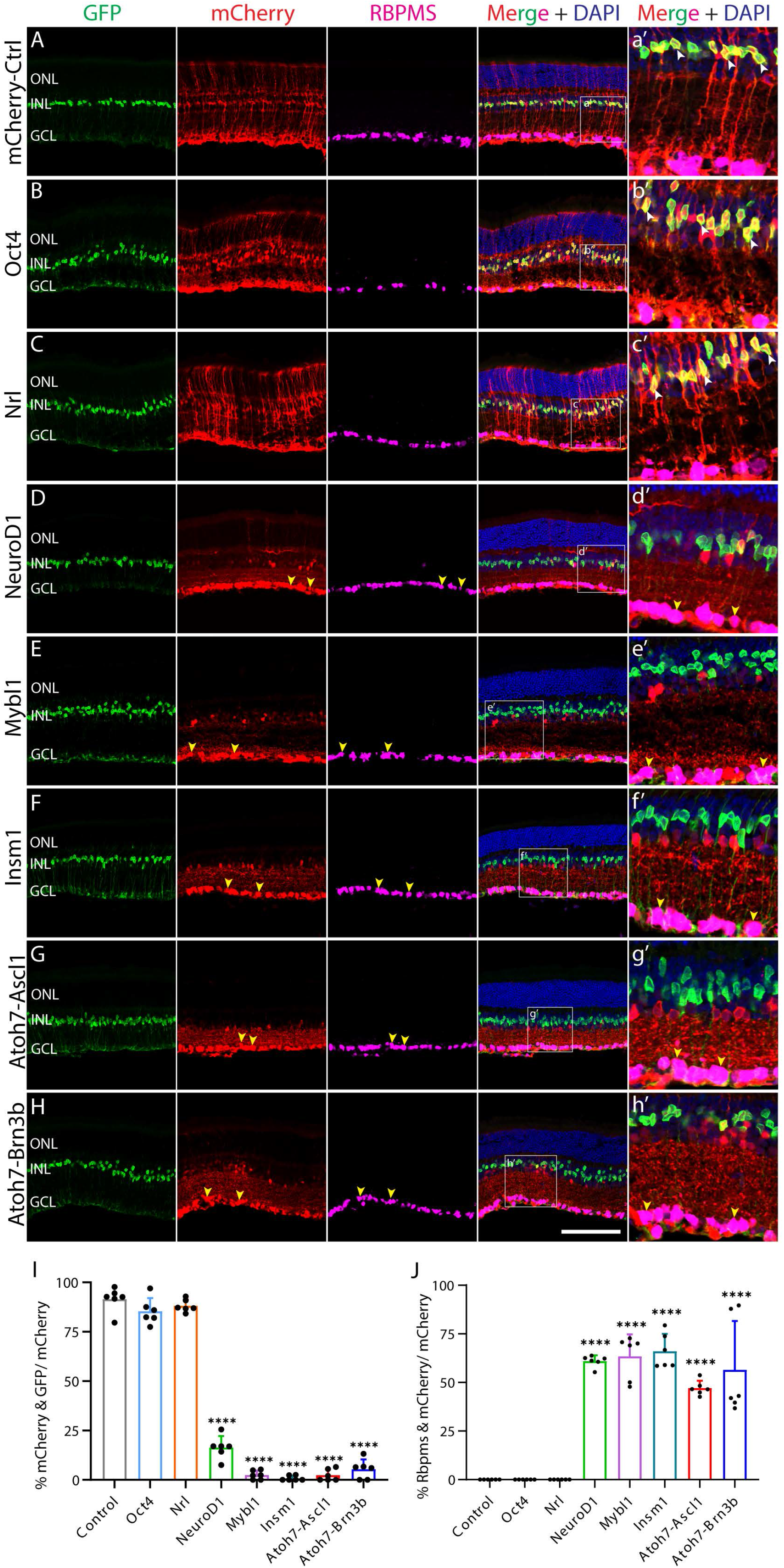
GFAP promoter constructs show insert-dependent ectopic mCherry expression in ganglion cells in adult mouse retina. **(A-H)** Representative immunostaining for GFP, mCherry and Rbpms expression in the retinas collected 21 days post GFAP AAV infection. **(a’-h’)** Outlined region shown at higher magnification. GFAP-mCherry **(A, a’)**, GFAP-Oct4-mCherry **(B, b’)** and GFAP-Nrl-mCherry **(C, c’)** showed almost complete co-localization of mCherry and Müller glia-specific Sun1-GFP. Arrows indicate co-labeled mCherry+/GFP+ cells. GFAP-NeuroD1-mCherry **(D, d’)**, GFAP-Mybl1-mCherry **(E, e’)**, GFAP-Insm1-mCherry **(F, f’)**, GFAP-Atoh7-Ascl1-mCherry **(G, g’)**, and GFAP-Atoh7-Brn3b-mCherry **(H, h’)** showed low mCherry+/GFP+ colocalization. High levels of mCherry expression in ganglion cells were observed in retinas infected with these constructs. Yellow arrowheads indicate Rbpms+/mCherry+ double positive cells. Quantification of mean percentage ± SD of mCherry+/GFP+ cells **(I)** and Rbpms+/mCherry+ cells **(J)**. Significance was determined via one-way ANOVA with Dunnett’s test: ***p < 0.001, ****p < 0.0001. Each data point in the bar graphs was calculated from an individual retina (*n* = 6). ONL, outer nuclear layer; INL, inner nuclear layer; GCL, ganglion cell layer. Scale bar = 100μm

**Figure 4:**
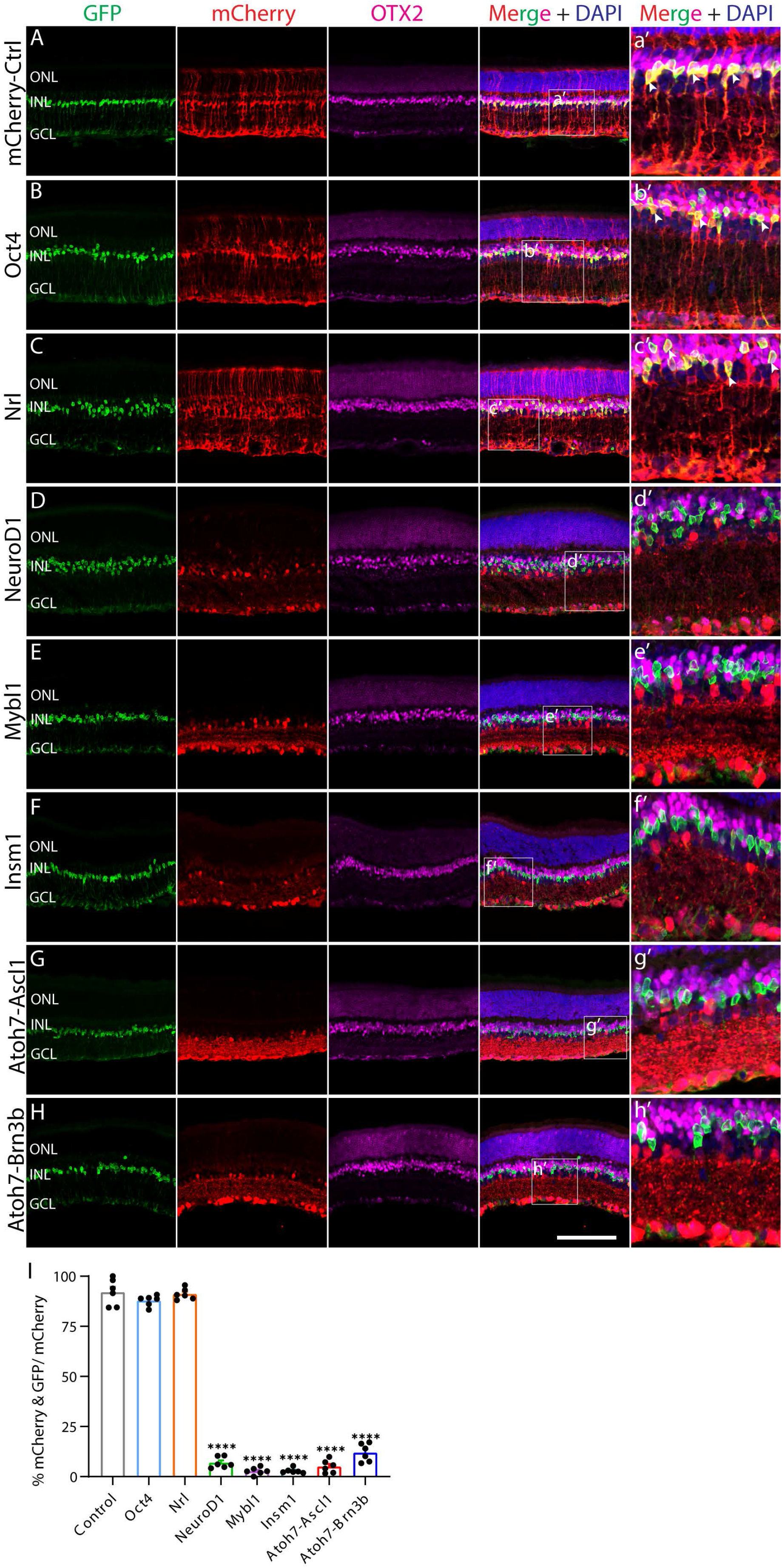
GFAP promoter constructs do not show insert-dependent ectopic mCherry expression in bipolar cells in adult mouse retina. **(A-H)** Representative immunostaining for GFP, mCherry and Otx2 expression in the retinas collected 21 days post GFAP AAV infection. **(a’-h’)** Outlined region shown at higher magnification. GFAP-mCherry **(A, a’)**, GFAP-Oct4-mCherry **(B, b’)** and GFAP-Nrl-mCherry **(C, c’)** showed almost complete co-localization of mCherry and Müller glia-specific Sun1-GFP. White arrowheads point to co-labeled mCherry+/GFP+ cells. GFAP-NeuroD1-mCherry **(D, d’)**, GFAP-Mybl1-mCherry **(E, e’)**, GFAP-Insm1-mCherry **(F, f’)**, GFAP-Atoh7-Ascl1-mCherry **(G, g’)**, and GFAP-Atoh7-Brn3b-mCherry **(H, h’)** showed low mCherry/GFP colocalization. No mCherry+/Otx2+ double positive cells were observed in retinas infected with these constructs. Quantification of mean percentage ± SD of mCherry+/GFP+ cells **(I)**. Significance was determined via one-way ANOVA with Dunnett’s test: ***p < 0.001, ****p < 0.0001. Each data point in the bar graphs was calculated from an individual retina (*n* = 6). ONL, outer nuclear layer; INL, inner nuclear layer; GCL, ganglion cell layer. Scale bar = 100μm

To further confirm that this observed ectopic mCherry expression was driven by the GFAP minipromoter, we retested several constructs that showed leaky neuronal expression using Cre-dependent Flexed AAV vectors (Schnütgen et al., 2003). To do this, we intravitreally injected 2 month old *GlastCreER;Sun1-GFP* mice with 7m8 AAV2 Cre-dependent flexed vectors expressing either mCherry alone, or mCherry in combination with Atoh7 and Brn3b, under the control of the ubiquitous EF1a promoter (Fig. 5A,B). Beginning four days following injection, we conducted four daily consecutive injections of 4-OHT to induce expression of both the flexed AAV construct and the transgenic Sun1-GFP construct. At 21 days following the fourth dose of 4-OHT injection, we then examined expression of both mCherry and Sun1-GFP, and observed selective Müller glia-specific expression of mCherry in both constructs tested indicated by extensive colocalization of GFP and mCherry expression (Fig. 5C,D). In contrast to previous reports (Xiao et al., 2021), we observe no evidence for transdifferentiation of Müller glia to retinal ganglion cells following misexpression of Atoh7 and Brn3b.

**Figure 5:**
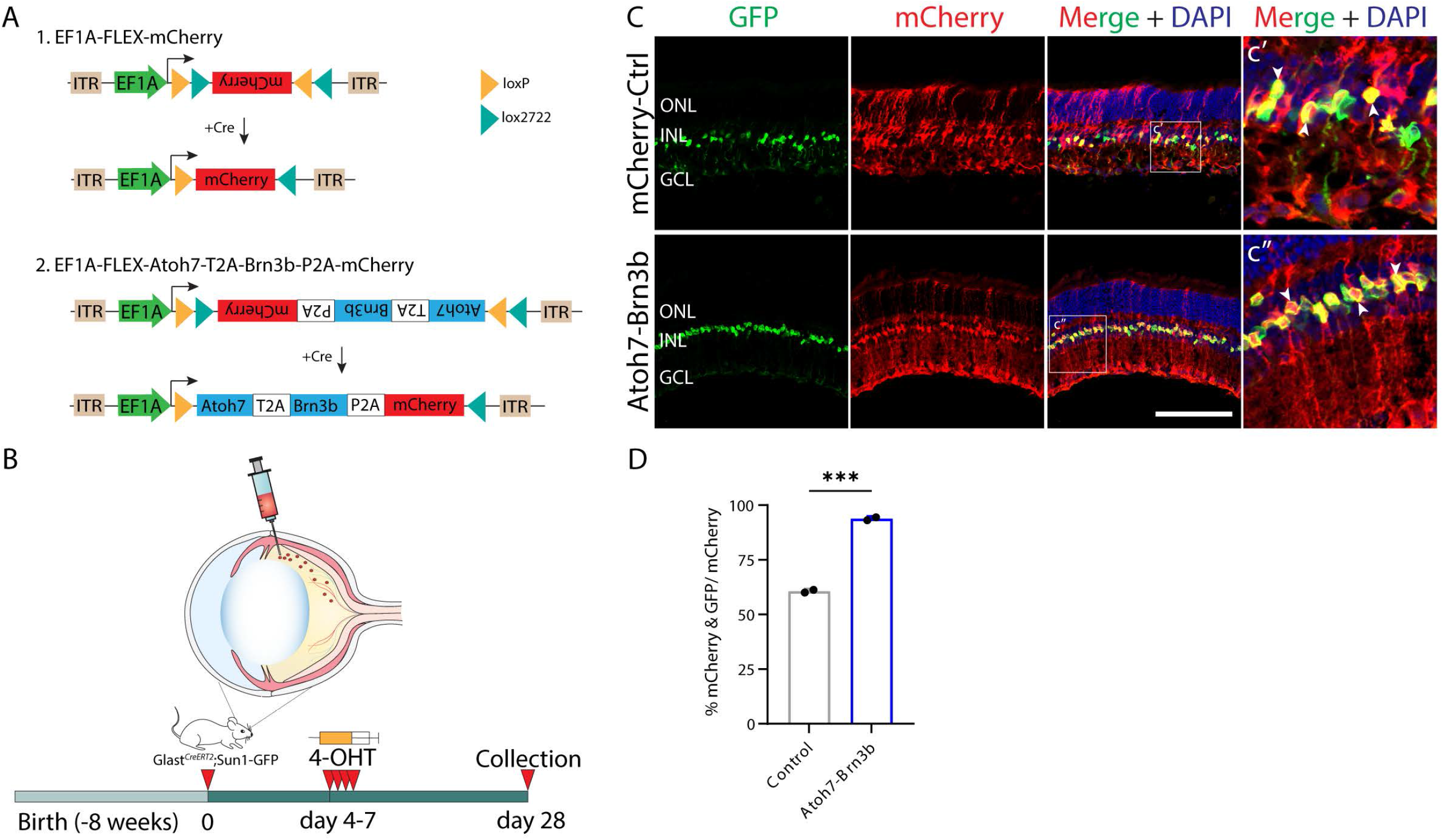
Inserts that show ectopic neuronal expression when expressed using the GFAP promoter show highly Müller glia-specific expression when expressing using GlastCreER-dependent EF1a FLEX AAV constructs. **(A)** A schematic of Cre-dependent FLEX-mCherry control and FLEX-Atoh7-Brn3b-mCherry constructs. **(B)** A schematic of the experimental workflow: two months old *GLASTCreERT2;Sun1-GFP* mice were intravitreally injected with FLEX AAV constructs and collected 21 days following the 4th dose of 4-OHT i.p injection for analysis. **(C)** Representative immunostaining for GFP and mCherry expression in FLEX AAV infected retinas collected 21 days post Cre-recombination induction. **(c’, c’’)** Outlined region shown at higher magnification. FLEX-Atoh7-Brn3b-mCherry showed extensive colocalization of GFP and mCherry. White arrowheads indicate mCherry+/GFP+ cells. **(D)** Quantification of mean percentage ± SD of mCherry+/GFP+ cells. Significance was determined via unpaired two-tailed t-test and Gaussian distribution: ***p < 0.001. Each data point in the bar graphs was calculated from an individual retina (*n* = 2). ONL, outer nuclear layer; INL, inner nuclear layer; GCL, ganglion cell layer Scale bar = 100μm.

## Discussion

In this study, we observed insert-dependent ectopic neuronal expression of GFAP minipromoter-driven mCherry expression for different AAV constructs. While GFAP-mCherry, the standard control used for experiments of this sort, consistently showed robust and selective glial-specific expression, the great majority of mCherry expression was restricted retinal amacrine and ganglion cells for 5 out of the 7 experimental overexpression constructs that were tested. This did not reflect Müller glia-to-neuron conversion, as these cells did not express Müller glia-derived Sun1-GFP, but instead implies that the insert sequences themselves dramatically altered the specificity of the GFAP minipromoter. This could reflect the action of cis-regulatory elements that bind transcription factors that promote neuronal-specific expression, the activity of the vector-encoded transcription factors on the GFAP minipromoter, or some combination of the two. Similar conclusions have been drawn following re-examination of claims of astrocyte-to-neuron conversion in brain (Chen et al., 2021; Hoang et al., 2021; Wang et al., 2021a), and these findings raise serious concerns about the accuracy of previously reported claims of AAV-mediated Müller glia-to-neuron conversion (Yao et al., 2018; Zhou et al., 2020; Xiao et al., 2021).

Since AAV vectors currently represent the least toxic and most effective method of gene delivery *in vivo*, this poses important challenges for future research into cell-based therapies. In mice, where transgenic reagents such as the *GlastCreER;Sun1-GFP* that can reliably and irreversibly label glia are readily available, prospective genetic lineage analysis should be performed routinely in conjunction with AAV-based manipulations. In this case, Cre-dependent FLEX-AAV constructs can also be used to further restrict the action of AAV-based constructs to glial cells and avoid ectopic neuronal expression. In species where genetic models are not readily available, or in preclinical studies in humans, this requires other approaches. These could include detailed morphological, physiological, and/or single cell RNA-Seq-based identification of reprogrammed glia and immature neurons.

## Acknowledgements

We thank I. Palazzo for comments on the manuscript. This study was supported by NIH National Eye Institute grants R01EY020560 and U01EY027267 and a Stein Innovation Award from Research to Prevent Blindness to S.B.

## Materials and Methods

### Mice

Mice were raised and housed in a climate-controlled pathogen-free facility on a 14/10 h light/dark cycle. Mice used in this study were *GlastCreERT2;Sun1-GFP*, which were generated by breeding. Maintenance and experimental procedures performed on mice were in accordance with the protocol approved by the Institutional Animal Care and USe Committee (IACUC) at the Johns Hopkins School of Medicine.

### GFAP-AAV Delivery

#### Intraperitoneal 4-Hydroxytamoxifen (4-OHT) Injection

To induce *Cre* recombination, *GLASTCreERT2;Sun1-GFP* mice at ∼3 weeks of age were intraperitoneally injected with 4-OHT (Sigma-Aldrich, #H6278-50mg) in corn oil (Sigma-Aldrich, #C8267-500ML) at 1mg/dose for three consecutive days.

#### Cloning, Production and Intravitreal Injection of Adeno-Associated Virus

The Addgene #50473 construct which contains a GFAP promoter was used in this study. The EGFP sequence was replaced by the mCherry sequence. Coding sequences of different transcription factors were synthesized by GeneWiz. AAV constructs were packaged into AAV2.7m8 by Boston Children’s Hospital Viral Core. Following 4-OHT i.p induction, two months old *GLASTCreERT2;Sun1-GFP* mice were intravitreally injected with GFAP AAV constructs using a microsyringe with a 33G blunt-ended needle. Titre and injection volume for each construct are listed below:

| AAV-GFAP Constructs | Titre (gc/ml) | Injection volume (μl) |
| --- | --- | --- |
| GFAP-mCherry | 1.96E<sup>13</sup> | 1μl/eye |
| GFAP-Oct4-P2A-mCherry | 2.58E<sup>13</sup> | 1μl/eye |
| GFAP-Nrl-P2A-mCherry | 2.58E<sup>13</sup> | 1μl/eye |
| GFAP-NeuroD1-P2A-mCherry | 2.83E<sup>13</sup> | 1μl/eye |
| GFAP-Mybl1-P2A-mCherry | 2.73E<sup>13</sup> | 1μl/eye |
| GFAP-Insm1-P2A-mCherry | 5.76E<sup>12</sup> | 1μl/eye |
| GFAP-Atoh7-T2A-Ascl1-P2A-mCherry | 7.60E<sup>12</sup> | 2μl/eye |
| GFAP-Atoh7-T2A-Brn3b-P2A-mCherry | 8.50E<sup>12</sup> | 2μl/eye |

### FLEX-AAV Delivery

Two months old *GLASTCreERT2;Sun1-GFP* mice were intravitreally injected with AAV-FLEX constructs using a microsyringe with a 33G blunt-ended needle. Four days following AAV transduction, mice were i.p induced with 4 daily doses of 4-OHT (1mg/dose in corn oil) and collected 21 days following the fourth dose for analysis. Titre and injection volume for each construct are listed below:

| AAV-FLEX Constructs | Titre (gc/ml) | Injection volume (μl) |
| --- | --- | --- |
| EF1a-DIO-mCherry (Addgene#50462) | 1.30E<sup>13</sup> | 1μl/eye |
| EF1a-DIO-Atoh7-T2A-Brn3b-P2A-mCherry | 6.70E<sup>12</sup> | 2μl/eye |

#### Fixation, Sectioning, Immunohistochemistry and Imaging

Collection and immunohistochemical analysis of retinas was performed as described previously (Hoang et al. 2020). Briefly, mouse eye globes were collected at 8 and 21 days post AAV infection and were fixed in 4% paraformaldehyde for 4 h at 4°C. Retinas were dissected in 1x PBS and incubated in 30% sucrose overnight at 4°C. Retinas were then embedded in OCT (VWR, #95057-838), cryosectioned at 16μm thickness, and stored at −20°C. Sections were dried for 30 mins in a 37°C incubator and washed 3×5 min with 0.1% TritonX-100 in PBS (PBST) and incubated in 10% Horse Serum (ThermoFisher, #26050070), 0.4% TritonX-100 in 1x PBS (blocking buffer) for 2 h at room temperature (RT). Sections were then incubated with primary antibodies in the blocking buffer overnight at 4°C. Primary antibodies used were Chicken anti-GFP (ThermoFisher, #A10262, 1:400), Rabbit anti-RFP (Abcam, #ab124754, 1:400), Goat anti-RFP (Rockland, #200-101-379, 1:400), Goat anti-Otx2 (R&D Systems, #AF1979, 1:200), Rabbit anti-Rbpms (ThermoFisher, #15187-1-AP, 1:400), Mouse anti-Tfap2a (Abnova, #H00007020-M01, 1:200), Rabbit anti-Sox9 (Sigma-Aldrich, #AB5535, 1:400).

Excess antibodies were removed by washing sections 4×5min with PBST before secondary antibodies incubation in the blocking buffer for 2 h at RT. Secondary antibodies used were Donkey anti-Chicken 488 (Sigma-Aldrich, #SAB4600031-250UL, 1:400), Donkey anti-Rabbit 568 (ThermoFisher, #A-10042, 1:400), Donkey anti-Goat 568(ThermoFisher, #A11057, 1:400), Donkey anti-Goat 633 (ThermoFisher, A-21082, 1:400), Donkey anti-Rabbit 647 (ThermoFisher, #A-31573, 1:400), Donkey anti-Mouse 647 (ThermoFisher, #A-31571, 1:400). Sections were then counterstained with DAPI in PBST, washed 4×5min in PBST and mounted with ProLong Gold Antifade Mountant with DAPI (Invitrogen, #P36935) under coverslips, air-dried, and stored at 4°C. Fluorescent images were captured using a Zeiss LSM 700 confocal microscope.

#### Cell Quantification and Statistical Analysis

Sox9+/GFP+, mCherry+/GFP+, Tfap2a+/mCherry+, Rbpms+/mCherry cells were counted and divided by the total number of mCherry+ cells from a random, fixed size region per retina. Each data point in the bar graphs was calculated from an individual retina. All cell quantification data were graphed and analyzed using GraphPad Prism 9. Analysis between 2 samples used unpaired two-tailed t-test and Gaussian distribution. One-way ANOVA with Dunnett’s test were used for analysis between 3 or more samples. All results are presented as mean ± SD.

