## Supplemental Figures 1-3 and legends for "Ectopic insert-dependent neuronal expression of GFAP promoter-driven AAV constructs in adult mouse retina"

**Supplemental figure legends:**

**Figure S1: Ectopic insert-dependent mCherry expression from GFAP promoter-derived AAV constructs in amacrine cells is evident at 8 days following transduction.**

**(A-D)** Representative immunostaining for GFP, mCherry and Tfap2a expression in the retinas collected 8 days post GFAP AAV infection. **(a'-d')** Outlined region shown at higher magnification. GFAP-mCherry **(A, a')** showed almost complete co-localization of construct-derived mCherry and Müller glia-specific Sun1-GFP. White arrowheads indicate mCherry+/GFP+ cells. GFAP-Insm1-mCherry **(B, b')**, GFAP-Atoh7-Ascl1-mCherry **(C', c')** and GFAP-Atoh7-Brn3b-mCherry **(D, d')** showed low mCherry/GFP colocalization. mCherry expression is observed in amacrine cells with these GFAP AAV constructs. Yellow arrowheads indicate Tfap2a+/mCherry+ cells. Quantification of mean percentage ± SD of mCherry+/GFP+ cells **(E)** and Tfap2a+/mCherry+ cells **(F)**. Significance was obtained via one-way ANOVA with Dunnett's test: $**p < 0.01$, $***p < 0.001$, $****p < 0.0001$. Each data point in the bar graphs was calculated from an individual retina ($n = 4$). ONL, outer nuclear layer; INL, inner nuclear layer; GCL, ganglion cell layer. Scale bar = 100μm.

**Figure S2: Ectopic insert-dependent mCherry expression from GFAP promoter-derived AAV constructs in retinal ganglion cells is evident at 8 days following transduction.**

**(A-D)** Representative immunostaining for GFP, mCherry and Rbpms expression in the retinas collected 8 days post GFAP AAV infection. **(a'-d')** Outlined region shown at higher magnification. GFAP-mCherry **(A, a')** showed almost complete co-localization of construct-derived mCherry and Müller glia-specific Sun1-GFP. White arrowheads indicate mCherry+/GFP+ cells. GFAP-Insm1-mCherry **(B, b')**, GFAP-Atoh7-Ascl1-mCherry **(C', c')** and GFAP-Atoh7-Brn3b-mCherry **(D, d')** showed low mCherry/GFP colocalization. Robust mCherry expression is observed in ganglion cells with these GFAP AAV constructs. Yellow arrowheads indicate Rbpms+/mCherry+ cells. Quantification of mean percentage ± SD of mCherry+/GFP+ cells **(E)** and Rbpms+/mCherry+ cells **(F)**. Significance was determined via one-way ANOVA with Dunnett's test: $****p < 0.0001$. Each data point in the bar graphs was calculated from an individual retina ($n = 4$). ONL, outer nuclear layer; INL, inner nuclear layer; GCL, ganglion cell layer. Scale bar = 100μm

**Figure S3: No ectopic insert-dependent mCherry expression from GFAP promoter-derived AAV constructs is detected in bipolar cells.**

**(A-D)** Representative immunostaining for GFP, mCherry and Otx2 expression in the retinas collected 8 days post GFAP AAV infection. **(a'-d')** Outlined region shown at higher magnification. GFAP-mCherry **(A, a')** showed almost complete co-localization of construct-derived mCherry and Müller glia-specific Sun1-GFP. White arrowheads point to co-labeled mCherry+/GFP+ cells. No mCherry+/GFP+ double positive or mCherry+/Otx2+ double positive cells were observed in retinas infected with GFAP-Insm1-mCherry **(B, b')**, GFAP-Atoh7-Ascl1-mCherry **(C', c')** and GFAP-Atoh7-Brn3b-mCherry **(D, d')**. ONL, outer nuclear layer; INL, inner nuclear layer; GCL, ganglion cell layer. Scale bar = 100μm

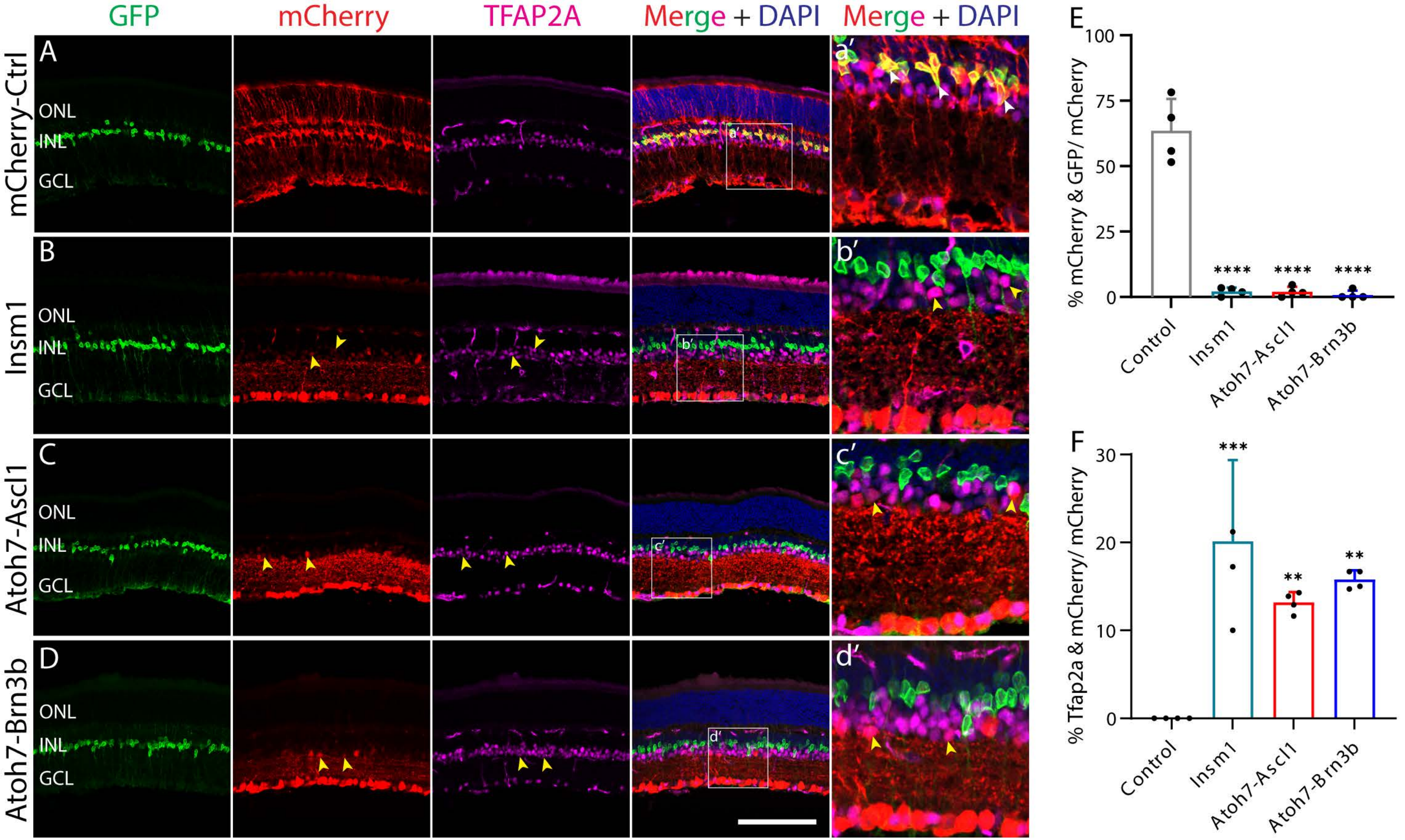

GFP
mCherry
TFAP2A
Merge + DAPI
Merge + DAPI
mCherry-Ctrl
Insm1
Atoh7-Ascl1
Atoh7-Brn3b
A
B
C
D
a′
b′
c′
d′
ONL
INL
GCL
E
% mCherry & GFP/ mCherry
100
75
50
25
0
****
****
****
Control
Insm1
Atoh7-Ascl1
Atoh7-Brn3b
F
% Tfap2a & mCherry/ mCherry
30
20
10
0
***
**
**
Control
Insm1
Atoh7-Ascl1
Atoh7-Brn3b


Fig. S1

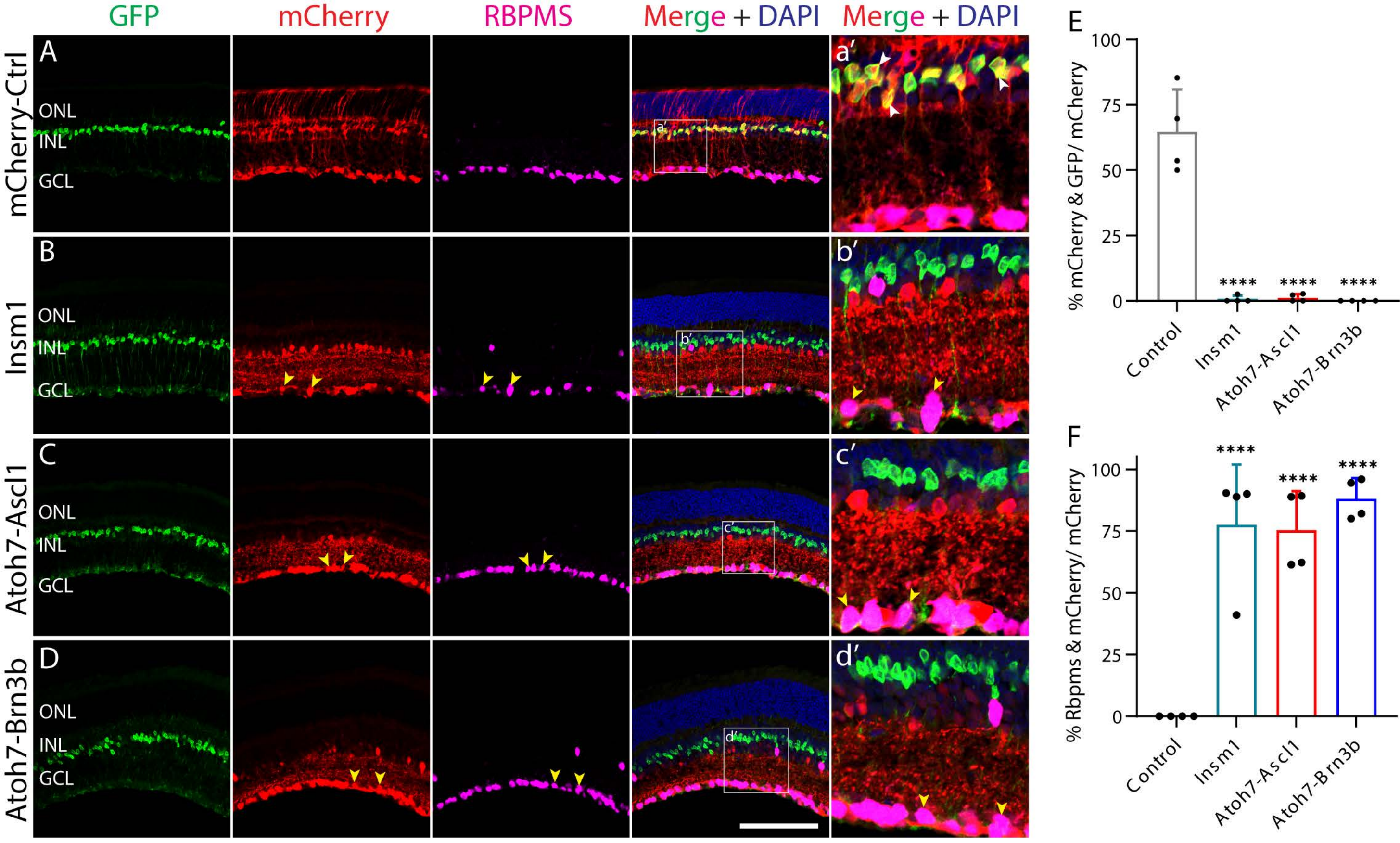
GFP
mCherry
RBPMS
Merge + DAPI
Merge + DAPI
A
B
C
D
a′
b′
c′
d′
mCherry-Ctrl
Insm1
Atoh7-Ascl1
Atoh7-Brn3b
ONL
INL
GCL
E
% mCherry & GFP/ mCherry
100
75
50
25
0
Control
Insm1
Atoh7-Ascl1
Atoh7-Brn3b
****
F
% Rbpms & mCherry/ mCherry

Fig. S2

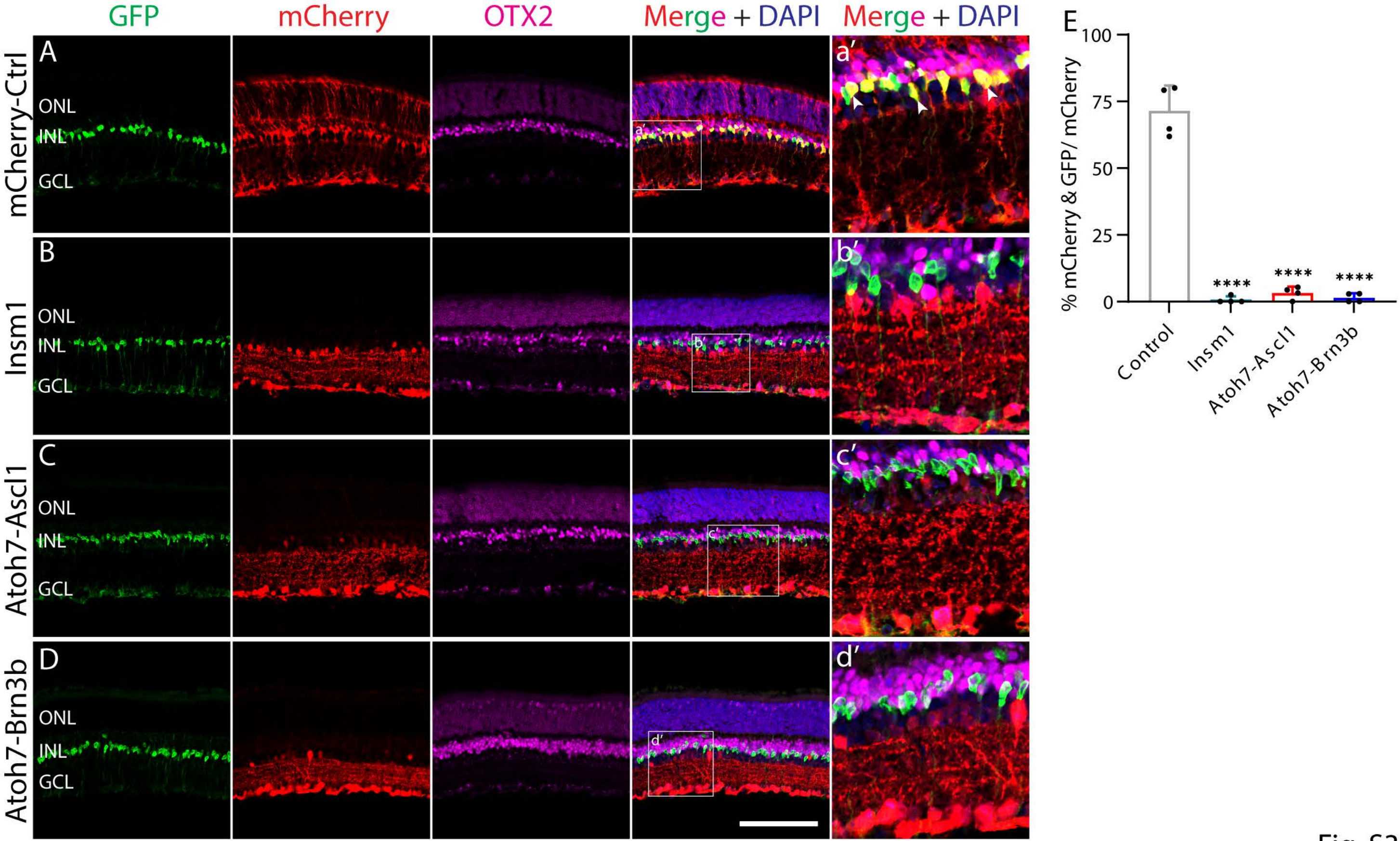
GFP
mCherry
OTX2
Merge + DAPI
Merge + DAPI
A
B
C
D
E
a′
b′
c′
d′
mCherry-Ctrl
Insm1
Atoh7-Ascl1
Atoh7-Brn3b
ONL
INL
GCL
% mCherry & GFP/ mCherry
100
75
50
25
0
****
Control
Insm1
Atoh7-Ascl1
Atoh7-Brn3b

Fig. S3
